# Profiling tumor immune microenvironment of non-small cell lung cancer using multiplex immunofluorescence

**DOI:** 10.1101/2021.05.28.446005

**Authors:** Haoxin Peng, Xiangrong Wu, Ran Zhong, Tao Yu, Xiuyu Cai, Jun Liu, Yaokai Wen, Yiyuan Ao, Jiana Chen, Yutian Li, Hongbo Zheng, Yanhui Chen, Zhenkui Pan, Jianxing He, Wenhua Liang

## Abstract

**Purpose:** We attempt to profile the tumor immune microenvironment (TIME) of non-small cell lung cancer (NSCLC) by multiplex immunofluorescence (MIF).

**Experimental Design:** MIF test was performed on 681 NSCLC cases in our center between 2009 and 2011. The number, density, proportion and correlation of 26 types of immune cells in tumor nest and tumor stroma were evaluated. An unsupervised consensus clustering approach was utilized to identify robust clusters of patients. Immune-related risk score (IRRS) was constructed for prognosis prediction for disease-free survival (DFS).

**Results:** The landscape of TIME was illustrated, revealing some close interactions particularly between intrastromal neutrophils and intratumoral regulatory T cells (Treg) (r^2^ = 0.439, P < 0.001), intrastromal CD4+CD38+ T cells and intrastromal CD20-positive B cells (r^2^ = 0.539, P < 0.001), and intratumoral CD8-positive T cells and intratumoral M2 macrophages expressing PD-L1 (r^2^ = 0.339, P < 0.001). Three immune subtypes correlated with distinct immune characteristics and clinical outcomes were identified. The immune-activated subtype had the longest DFS and demonstrated the highest infiltration of CD4-positive T cells and CD20-positive B cells. The immune-defected subtype had the highest levels of cancer stem cells and macrophages. The immune-exempted subtype had the highest levels of neutrophils and Treg. The IRRS based on six robust prognostic biomarkers showed potential ability for risk stratification (high vs. median vs. low) and prediction of five-year DFS rates (43.1% vs. 37.9% vs. 23.2%, P<0.001).

**Conclusions:** Our study profiled the intricate and intrinsic structure of TIME in NSCLC, which showed potency in subtyping and prognostication.

**Translational Relevance:** Significant unmet need exists in understanding the tumor immune microenvironment (TIME) of non-small cell lung cancer (NSCLC) and its correlation with prognosis. In this retrospective cohort study (n = 681), we profiled the immune landscape of NSCLC in situ and identified a novel stratification of TIME by three immune subtypes: immune-activated, immune-exempted, and immune-defected using multiplex immunofluorescence for testing 26 kinds of immune cells. Each of the immune subtypes was correlated with distinct composition, spatial distribution, and functional orientation of immune cells, and accordingly indicating significantly different disease-free survival (DFS). Close interactions were observed for several kinds of immune cells, including neutrophils and regulatory T cells, CD4+CD38+ T cells and CD20-positive B cells, and CD8-positive T cells and M2 macrophages. We also developed the immune-related risk score (IRRS) with different immune characteristics based on six robust immune biomarkers in TIME and evaluated its role in risk stratification and prognosis prediction of DFS. This study might bring potential clinical implementations for the design of novel immunotherapies and the optimization of combined strategies.

## Introduction

According to the global cancer statistics reported by the International Agency for Research on Cancer, lung cancer (LC) is the second-most commonly diagnosed cancer and the most common cause of cancer death worldwide (1). Non-small cell lung cancer (NSCLC) accounts for around 85% of LC, and it encompasses two major histological subtypes, lung adenocarcinoma, and squamous cell lung cancer, respectively (2). Despite the immense improvements in new drugs and systemic therapy, the five-year overall survival (OS) rate for advanced NSCLC patients was less than 5% (3).

Emerging evidence shows that the immune microenvironment (TIME) is the key determinant of LC development and patients’ prognosis (4,5). The TIME mainly contains neoplastic cells, stromal cells, and diverse immune cells, and these components interact mutually through complex cellular and molecular mechanisms, which influence tumor progression, metastasis, and clinical outcomes like treatment tolerance (6). The location, type, density, and functional state of immune cells constitute the TIME’s immune contexture, varying in patients with NSCLC. The immune cells may have dual impacts for both anti-tumor and pro-tumor effects. For instance, CD8+ T cells, NK cells mediate antitumoral responses, demonstrating a better OS, disease-free survival (DFS), and PFS. On the contrary, the regulatory T cells (Tregs) can secrete inhibitory cytokines, such as transforming growth factor (TGF)-β and interleukin (IL)-10, contributing to LC progression via angiogenesis, immunosuppression through inhibition of the anti-tumor effect of T-helper (Th1) cells as well as attracting activated Th2 cells (7-10).

Immunotherapy, mainly enhancing the anti-tumor immune responses through targeting T cell regulatory pathway in TIME, has shown enormous potential and promising results for improving disease control in NSCLC patients in recent years (11). For instance, the five-year OS rate triggered by immunotherapy, especially immune checkpoint blockades (ICB), now surpasses 25% for patients with high PD-L1 expression (tumor proportion score≥50%) (12). But the long-term clinical benefits occur only in a limited portion of patients (13). Current studies pointed that PD-L1, tumor mutational burden, and intratumoral heterogeneity may provide hints of prognosis with immunotherapy in NSCLC. However, there is no consensus regarding the best predictive biomarker of prognosis (14). Still, we know little about NSCLC TIME and how this information could be utilized to design appropriate therapies for distinct patient subgroups. Hence, a vital unmet need is to investigate the critical components and related cellular and molecular mechanisms responsible for immune responses, exhaustion, or ignorance to modify the TIME and design effective therapies.

Previous traditional immunohistochemistry (IHC)-based studies are usually limited to a relatively small sample size and few immune cell types, making it insufficient to exhibit the immune landscape of TIME fully. Recently, the multiplex immunofluorescence (MIF) approach has been demonstrated to provide a unique perspective into the spatial relationships among immune cells, stromal cells and tumor cells within the complex TIME. The MIF also avoids the traditional shortcomings of IHC, such as the low reproducibility and the subjective scoring system (15,16).

In this study, we described the immune landscape of NSCLC in situ and identified a novel stratification of TIME by three immune subtypes using MIF. We also established the IRRS as a robust prognostic biomarker for DFS.

## Materials and Methods

### Patient cohort

From 2009 to 2011, we collected a consecutive series of 681 NSCLC patients from stage I to III operated lobectomy/sub-lobectomy and lymph node dissection at the First Affiliated Hospital of Guangzhou Medical University. Written informed consent was obtained from all patients for permitting MIF analyses of biological samples. The study was conducted following the Declaration of Helsinki (as revised in 2013) (17).

Inclusive criteria were: (1) single primary NSCLC; (2) stage I to III; (3) underwent anatomical resection in combination with lymphadenectomy (systematic lymph node sampling or systematic lymph node dissection) according to the National Comprehensive Cancer Network (NCCN) criteria (18,19); (4) all resected tissues and lymph nodes were confirmed by pathology finally; and (5) sufficient resected tissues for MIF test. Patients were excluded if any of the following criteria meet: (1) multiple LC; (2) small cell lung cancer (SCLC) or non-invasive LC like LUAD in situ and minimally invasive LUAD; (3) diagnostic biopsy in pre-operation; and (4) preoperative neoadjuvant therapy.

### Multiplex immunofluorescence detection

The multiplexed immunofluorescence staining was conducted at Genecast Biotechnology Co., Ltd. (Beijing, China). Briefly, 4μm thick section was cut from FFPE(formalin-fixed paraffin-embedded) lung cancer tissues for each panel detection. The slides were deparaffinized, rehydrated, and subjected to epitope retrieval by boiling in Tris-EDTA buffer (pH=9; Klinipath #643901, Duiven, the Netherlands) for 20 minutes at 97°C. Endogenous peroxidase was then blocked by incubation in Antibody Diluent/Block (PerkinElmer #72424205, Massachusetts, USA) for 10 minutes, and protein was subsequently blocked in 0.05% Tween solution containing 0.3% bovine serum albumin for 30 minutes at room temperature. Only one antigen was detected in each round, including primary antibody incubation, secondary antibody incubation, tyramine signal amplification(TSA) visualization, followed by labeling the next antibody after epitope retrieval and protein blocking as before. CD4, CD20, CD38, CD66b and FOXP3 for panel 1, and CD8, CD68, CD133, CD163, and programmed cell death protein ligand-1 (PD-L1) for panel 2 were sequentially detected.

Primary antibodies for CD8 antibody (ZA-0508, clone SP16, Zsbio, 1:100), CD20 (ab9475, abcam, 1:50), Zsbio, 1:100), CD38 (ZM0422, clone SPC32, Zsbio, 1:400), CD66b (ab214175, polyclonal antibody, abcam, 1:50), CD68 (ZM-0060, clone KP1, Zsbio, 1:100), CD163 (ZM-0428, clone 10D6, PD-L1 (13684s, clone E1L3N, CST, 1:100), FoxP3 (ab20034, clone 236A/E7, abcam, 1:100) were incubated for 1 h at room temperature, CD4 (ZM0418, clone UMAB64, Zsbio, 1:200) and CD133 (ab19898, polyclonal antibody, abcam, 1:400) were incubated for overnight at 4□.

Anti-rabbit/mouse horseradish peroxidase (HRP) antibodies (Zsbio # PV-6002 or PV-8000) were used as the secondary antibody and incubated at 37□ for 10 min. TSA visualization was then performed with the opal seven-color multiplex immunohistochemistry Kit (NEL797B001KT, PerkinElmer, Massachusetts, USA), containing fluorophores (DAPI), Opal 520 (CD20 and CD163), Opal 540 (CD38), Opal 570 (PD-L1 and CD4), Opal 620 (CD8), Opal 650 (CD66b and CD133), Opal 690 (CD68 and FoxP3) and TSA Coumarin system (NEL703001KT, PerkinElmer, Massachusetts, USA). After labeling all of the 5 antigens for each panel, microwave treatment (MWT) was performed to remove the TSA-antibody complex with Tris-EDTA buffer (pH=9; Klinipath #643901, Duiven, the Netherlands) for 20 minutes at 97°C. All the slides were counterstained with 4’,6-Diamidino-2-Phenylindole (DAPI) for 5 min and were enclosed in Antifade Mounting Medium (NobleRyder #I0052, Beijing, China), prepared for imaging. Fresh whole-tissue section cuts from normal human tonsils were included in each staining batch as positive control and assessed the interexperimental reproducibility.

Slides were scanned using the PerkinElmer Vectra (Vectra 3.0.5; PerkinElmer, Massachusetts, USA). Multispectral images were unmixed with spectral libraries built from single stained tissue images for each antigen, using the inForm Advanced Image Analysis software (inForm 2.3.0; PerkinElmer, Massachusetts, USA).

For batch analysis, an algorithm is acquired by training 10 to 15 representative multispectral images. Then tissue segmentation and cell segmentation were conducted with the algorithm. An experienced pathologist determined appropriate positive threshold X for each biomarker. We defined X, 2X, 3X as the threshold of the signal intensity of 1+, 2+, 3+ respectively, and the POS value equals X+2X+3X. Both the density (n/mm^2^), number (n/sight), and percentage (%/sight) of immune markers in tumor nest (TN) and tumor stroma (TS) were calculated. Histochemistry score (H-score) was analyzed with the formula of H-score = (3+)%×3+ (2+)%×2+ (1+)%×1. In total, 26 kinds of immune cells, including 66 kinds of immune biomarkers, were test and calculated. Immune cell types represented by biomarkers were labeled through literature retrieval (20) (Supplementary Table S1).

### Defining the immune landscape

The immune landscape of NSCLC was conducted with a MIF test for 681 cases, demonstrating the intricate and intrinsic structure of TIME and visualizing the distribution of individual patients.

### Discovery of the immune subtypes

Unsupervised consensus clustering is a class discovery approach to detect unknown possible clusters consisting of individual items with similar intrinsic features (21). Based on the proportion of 26 kinds of immune cells both in TN and TS, distinct subgroups of 681 samples were identified, during which 80% of the samples were extracted 100 times in turn and a hierarchical clustering analysis was performed based on the Euclidean distance between data points. The consensus clustering results were subsequently tested using the cumulative distribution function (CDF) plot corresponding to the consensus matrices. Then, the results of clustering were verified employing principal component analysis (PCA).

### Evaluating the cellular and clinical characteristics correlated with the immune subtypes

Kruskal-Wallis (K-W) test and boxplots were used to visualize the disparities of immune cell proportion among different clusters. The log-rank test was initially employed to evaluate the prognostic significance of immune subtypes. The multivariable Cox proportional hazards regression analysis was then used for further assessment with adjustments for age, sex, T stage, N stage, vascular cancer embolus, and number of lymph nodes resection, and DFS was considered as the endpoint. We also utilized the chi-square test to investigate the heterogeneity of clinical characteristics in three clusters.

### Profiling the prognostic value of immune biomarkers

Differences in the proportion of 26 kinds of immune cells among T stage, N stage, and clinical stage were analyzed through T-test. Multivariable Cox regression with age, sex, histological types, T stage, and N stage as covariates was utilized to identify the prognostic value of 66 immune biomarkers. We further classified the values of immune biomarkers into high-value and low-value subtypes by the optimal cut-off point according to the built-in risk scoring formula in X-tile and assessed the differences in DFS.

### Construction and validation of the immune-related risk score

The entire cohort (n = 681) was divided into the training cohort (n = 477) and the testing cohort (n = 204). Immune cells significantly associated with DFS through multivariable Cox regression (P<0.05) in the entire cohort were selected as the candidate factors. The LASSO Cox regression model was then used for profiling the most robust prognostic immune cells among the candidate factors, and the optimal lambda value was determined by 10-fold cross-validation (22). IRRS was ultimately conducted by the regression coefficients originated from multivariable Cox regression method to multiply the proportion of immune cells in the training cohort:

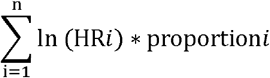

In which HR_*i*_ is the HR and proportion_*i*_ is the proportion for the ith immune cells. The multivariable Cox proportional hazards regression analysis with adjustments for sex, age, T stage, N stage, number of lymph node resection, and vascular cancer embolus was conducted to investigate the prognostic significance of IRRS in the training cohort. We further divided IRRS into high-risk, median-risk, and low-risk groups by the optimal truncation values to seek out the difference in DFS. The performance and robustness of IRRS in the training cohort was further tested in both the testing cohort and the entire cohort with the same formula.

### Statistical analysis

Using R package psych (version 2.0.12), the Spearman test was conducted to explore the correlations between immune cells in TN and TS, during which correlation coefficients and their P-values were calculated, and the correlations were shown in dot-line charts based on R package ggpubr (version 0.4.0). R package ConsencusClusterPlus (version 1.54.0) was used to perform unsupervised consensus clustering analysis to explore the intricate relationships of the immune cells, and the clustering results were verified with principal component analysis (PCA) using the R package FactoMineR (Version 2.4). Besides, we explored the differences of clinical characteristics among each cluster by percentage component bar chart and Sankey plot analysis, which were performed by R package ggplot2 (version 3.3.3) and ggalluvial (version 0.12.3), respectively. After that, in order to construct an IRRS-based prognostic model, we utilized LASSO regression using the glment package (version 4.0.2) in R software for high-dimensional data to select the most useful prognostic factors. The receiver operating characteristic (ROC) curve and time-dependent AUC curve was to test the accuracy of the IRRS model using R package timeROC (version 0.4) (23). We also performed multi-variable Cox regression analysis to assess whether IRRS was independent of other clinical characteristics. The survival distribution of DFS curves were estimated by Kaplan-Meier method and the two-sided log-rank test as implemented in the R package survminer (version 0.4.8). All statistical analyses were performed using R software (version 4.0.3). Data analysis was performed in SPSS software (version 23.0), and we used multi-variable Cox regression analysis to determine whether immune markers were independent of other clinical characteristics and significantly related to DFS. Hazard ratio (HR), 95% confidence interval (CI), and P-value for each immune marker were calculated. Pearson’s chi-square test and Fisher’s exact test were applied for comparison between categorical variables. Non-parametric analysis (Mann-Whitney U test or K-W test) was used for non-normally distributed rank/ordered variables and data, while continuous variables were analyzed by T-test. X-tile software was used to divide the values into several groups through the built-in risk scoring formula based on the combined model with the optimal cut-off points (24). All the P-values were two-sided, and the P-value <0.05 was considered statistically significant.

## Results

### Patients’ characteristics

Six hundred and eighty-one patients met the criteria and were included in this study, the baseline characteristics of whom were represented in Supplementary Table S2. Of the included cases, 321 (47.1%) were more than 60 years old and 398 (58.4%) were male. LUAD was the dominant histological subtype (479, 70.5%). Clinical stage IA, IIA, IB, IIB, IIIA, IIIB accounted for 22.0%, 31.1%, 16.2%, 5.5%, 24.8%, 0.5%, respectively.

### Immune landscape and interactions among immune cells

We exhibited the representative MIF images of each immune biomarker in TN and TS (Fig. 1A). The immune landscape of NSCLC both in TN and TS was demonstrated (Fig. 1B). Strong interactions were observed particularly on intrastromal CD20-positive B cell and intrastromal CD4+CD38+ T cell (r^2^ = 0.539, P = 9.23E-38), neutrophil and FOXP3-positive cell (r^2^ = 0.501, P = 5.17E-32 in TS and r^2^ = 0.552, P = 7.00E-40 in TN, respectively) as well as intrastromal neutrophils and intratumoral regulatory T cells (Treg) (r^2^ = 0.439, P = 9.48E-28), intratumoral CD38-positive T cell and intratumoral CD20-positive B cell (r^2^ = 0.525, P = 1.37E-35), intratumoral CD8-positive T cells and intratumoral M2 macrophages expressing PD-L1 (r^2^ = 0.339, P = 1.51E-16), intratumoral PD-L1-positive cell and intratumoral CD8-positive T cell (r^2^ = 0.407, P = 1.02E-20) and so on(Fig. 1 C-H, Supplementary Fig. S1, Supplementary Table S3 Supplementary Table S4). Moreover, strong correlations were also found between tumoral CD133-positive cell and tumoral M1 macrophage (r^2^ = 0.416, P = 9.14E-22) and M1 Macrophage without expressing PD-L1 (r^2^ = 0.451, P = 2.25E-29), more specifically. Given that CD133 is usually defined as a marker of cancer stem cell (CSC) of LC, the interaction between CSC and macrophage may contribute to the mechanisms underlying immune escape (25-28). Previous studies have deemed CD38 as the marker of activated CD4+ T cell and CD133 as the marker of CD8+ T cell stemness, and moderate correlation between CD4+CD38+ T cell and CD8+CD133+ T cell was observed (r^2^ = 0.316, P = 1.89E-14) in TS rather than in TN (29-32). In addition, moderate correlations for intratumoral CD68+PD-L1+ macrophage and intratumoral CD8+ T cell (r^2^ = 0.365, P = 5.82E-16), intratumoral M2 macrophage expressing PD-L1 and intratumoral CD8+ T cell (r^2^ = 0.339, P = 1.51E-16) was also presented, implying the intratumoral macrophage may play a role in mediating exhausted CD8-specific immune response (33). As for neutrophil and FOXP3-positive cell, previous studies have reported that tumor-associated neutrophil (TAN) can recruit FOXP3-positive cell through chemokine ligand 2 (CCL)-chemokine receptor-2 (CCR) and CCL17-CCR4 pathways to form immunosuppressive TIME and promote tumor progression (34). Moreover, we observed that the infiltrating levels of several cell types (neutrophils, CD68-positive macrophages, CD133-positive cells, M1 macrophages, CD68-positive macrophages expressing PD-L1, M2 macrophages without expressing PD-L1) were higher in TN than in TS (Supplementary Fig. S2).

**Figure 1.**
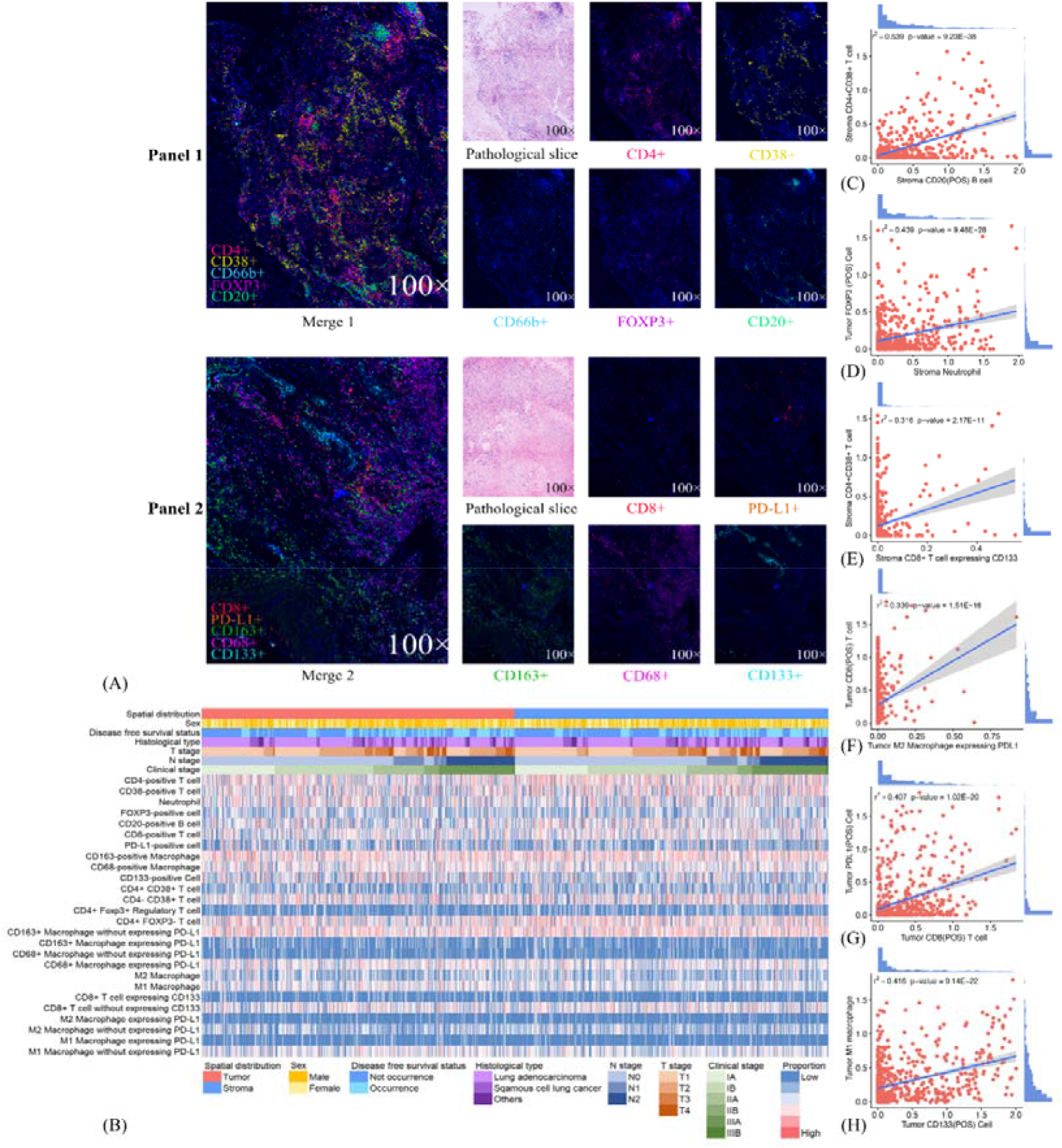
Ten immune biomarkers expression profile in resected lung cancer tissues. Single and merged immunofluorescence images and pathological slices are shown accordingly (A). Immune landscape of the TIME illustrates the log percentage (lg%) of each type of immune cell within tumor nest and tumor stroma. Each value corresponds to patients’ clinical characteristics, including sex, disease-free survival status, histological type, clinical stage, T stage, and N stage (B). The dotted line graphs illustrate the correlations between immune cells in TIME, and the bar graph shows the distribution of logarithmic percentage (lg%) of the proportion: (C) intrastromal CD20-positive B cells and intrastromal CD4+CD38+T cells; (D) intrastromal neutrophils and intratumoral FOXP3-positive cells; (E) intratumoral CD8-positive T cells and M2 macrophages expressing PD-L1; (F) intratumoral CD38-positive T cells and intratumoral CD20-positive B cells; (G) intratumoral CD8-positive T cells and intratumoral PD-L1-positive cells; (H) intratumoral CD133-positive cells and intratumoral M1 microphages.

### Components and clinical features of immune subtypes

Through conducting an unsupervised consensus clustering approach of 681 NSCLC cases, including cluster-consensus and item-consensus analyses, we identified three robust immune subtypes based on the proportion of immune cells in TIME (Fig. 2A). Kaplan-Meier survival analysis suggested significant differences in DFS among three immune subtypes, and subtype 1 had the longest DFS while subtype 3 showed the worst (HR 1.50, 95%CI 1.07-2.11, P = 0.019) (Fig. 2B), which were further supported by multivariable Cox regression analysis (HR 1.51 95%CI 1.05-2.17, P = 0.026). We also observed a trend for longer DFS in immune class 2 (HR 0.76, 95%CI 0.48-1.20, P = 0.237) compared with class 3. Patients in immune subtype 1 had a marginally better DFS than immune subtype 2 (P = 0.471) (Supplementary Table S5)

**Figure 2.**
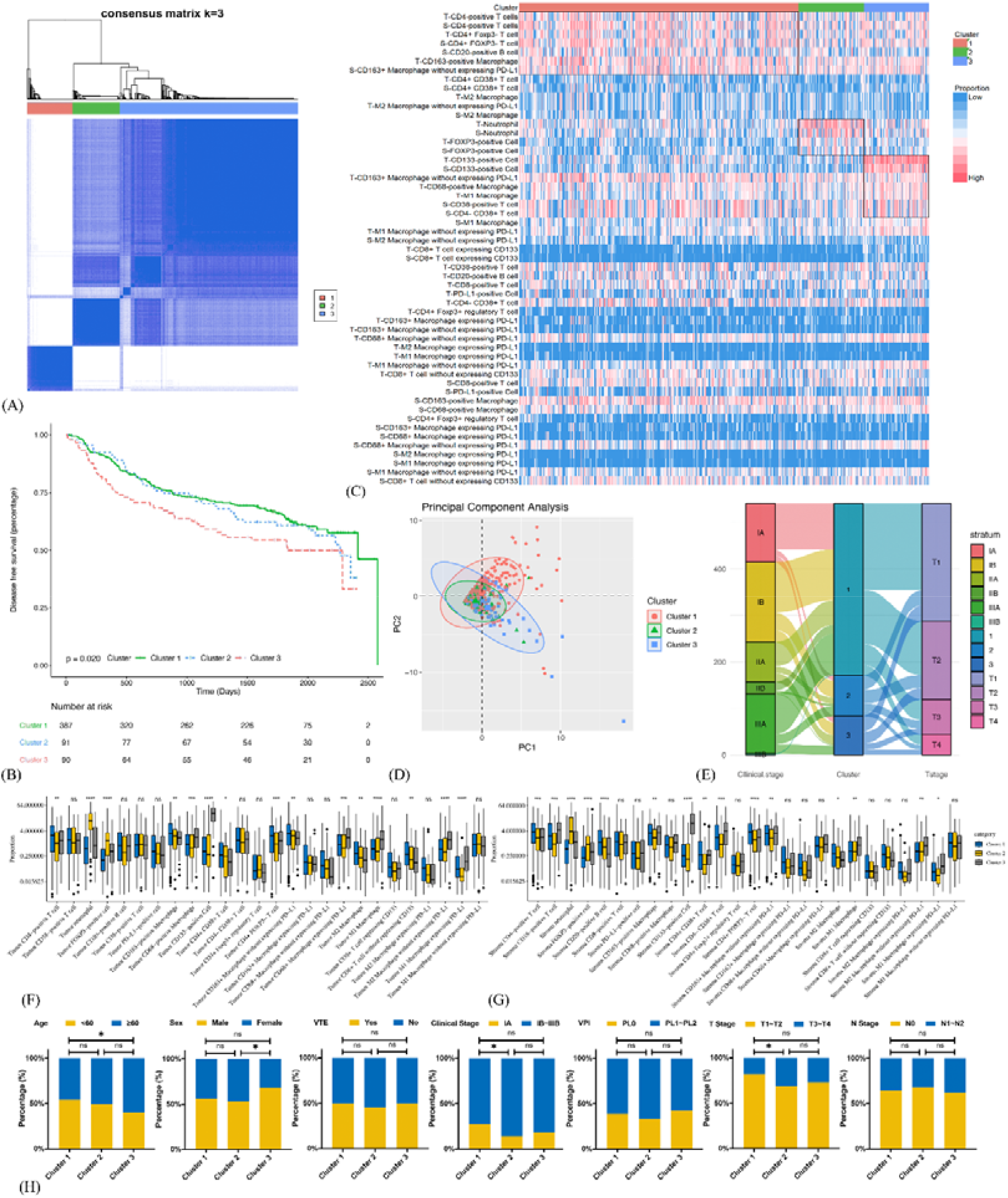
Identification and analysis of three clusters of 681 non-small cell lung cancer cases. (a) Consensus clustering matrix for k = 3. (b) Kaplan-Meier curves of three clusters. (c) Immune landscape of three clusters illustrates distinct cellular characteristics. (e) Principal component analysis of three clusters. (f) Sankey plots of three clusters. (g) Infiltration disparities of three clusters in tumor nest. (h) Infiltration disparities of three clusters in tumor stroma. (i) Chi-Square test reveals disparities of patients’ clinical characteristics among three clusters.

Distinct cellular features among three immune subtypes were shown (Fig. 2C-G). Immune subtype 1 accounted for 68.2% of enrolled patients, in which highest infiltration of CD4-positive T cells and CD20-positive B cells were observed and the proportion of both cells were higher in TS than TN (P < 0.001),. More specifically, CD4+CD38+ T cells and CD4+FOXP3-T cells were the major subsets of CD4-positive T cells rather than CD4+FOXP3+ regulatory T cells. Besides, the highest infiltrating levels of CD8-positive T cells were found in TN rather than in TS, indicating the active anti-tumor immunity. It has been reported that B cells organized in tertiary lymphoid structures (TLS) could present tumor antigens for activating CD4-positive cells, and B cells could also proliferate and differentiate into plasma cells for generating antibodies for antineoplastic effects with the help of IL-4 secreted by CD4-positive cells (35). Hence, CD4-positive T cells and CD20-positive B cells possibly act as the “guides” in participating anti-tumor responses indirectly in TS through secreting IFN-γ, recruiting and activating T cells, B cells, and NK cells, while CD8-positive T cells tend to directly kill tumor cells in TN. Thus this subtype was assumed to be “immune-activated” (Fig. 3). Moreover, we also found relatively a high infiltration of M2 macrophages in this subtype. Considering that M2 macrophage was usually associated with pro-tumor effects like angiogenesis and immunosuppression, it suggested the existence of intracluster heterogeneity (36). Further investigation for the functional state showed that the majority was the M2 macrophages without expressing PD-L1 rather than expressing PD-L1, implying the immunosuppressive function had not yet been developed and the anti-tumor effects might still be dominant, which was consistent with the recent report (37). Immune subtype 3 accounted for 16.0% of the included patients, similar to immune subtype 2 (15.8%), which was characterized by the highest proportion of CD133-positive cells, M1 macrophages expressing PD-L1 and M2 macrophages without expressing PD-L1. CD133-positive cell, mainly CSC, could show unlimited capacity for self-renewal, which plays a vital role in inducing tumor recurrence, metastasis, and heterogeneous tumor cells (38). Previous studies have demonstrated an intimate connection between CSC and macrophage. For instance, CSC can recruit Tregs into TIME, which subsequently secret IL-10, TGF-β in mediating immunosuppressive microenvironment and induce macrophage to polarize into M2 subset, also known as the tumor-associated macrophage (TAM) (39). The TAM would also in turn impact CSC, like inducing epithelial-mesenchymal transition of CSC to promote tumor invasion (40). Consequently, macrophage can be “educated” to develop pro-tumor effects under the impact of CSC, and so this subtype was regarded as “immune-defected” (Fig. 3). Patients in immune-defected subtype were also associated with older age and a higher proportion of male (Fig. 2H). The highest infiltration of neutrophils and FOXP3-positive cells in TN and TS was found in immune subtype 2, and as mentioned above, we also observed strong correlation between neutrophils and FOXP3-positive cells. FOXP3-positive cells, mainly Tregs, are generally thought to disrupt anti-tumor immunity (41). Nevertheless, the prognostic effects of neutrophils in NSCLC are still conflicting to date. In general, high levels of N1 neutrophils showed superior outcomes while N2 mainly indicated the negative, possibly through releasing matrix metallopeptidases-9 (MMP) and elastase to drive the metastasis of LC cells (42,43). Moreover, the levels of other infiltrating immune cells were the lowest in this subtype, and the T stage and clinical stage were more advanced (P < 0.05) (Fig. 2H). Therefore, the formation of an immunosuppressive microenvironment and lack of immune responses made cancer cells have the privileges and immunities from immune attack and so was regarded as “immune-exempted” (Fig. 3) (44).

**Figure 3.**
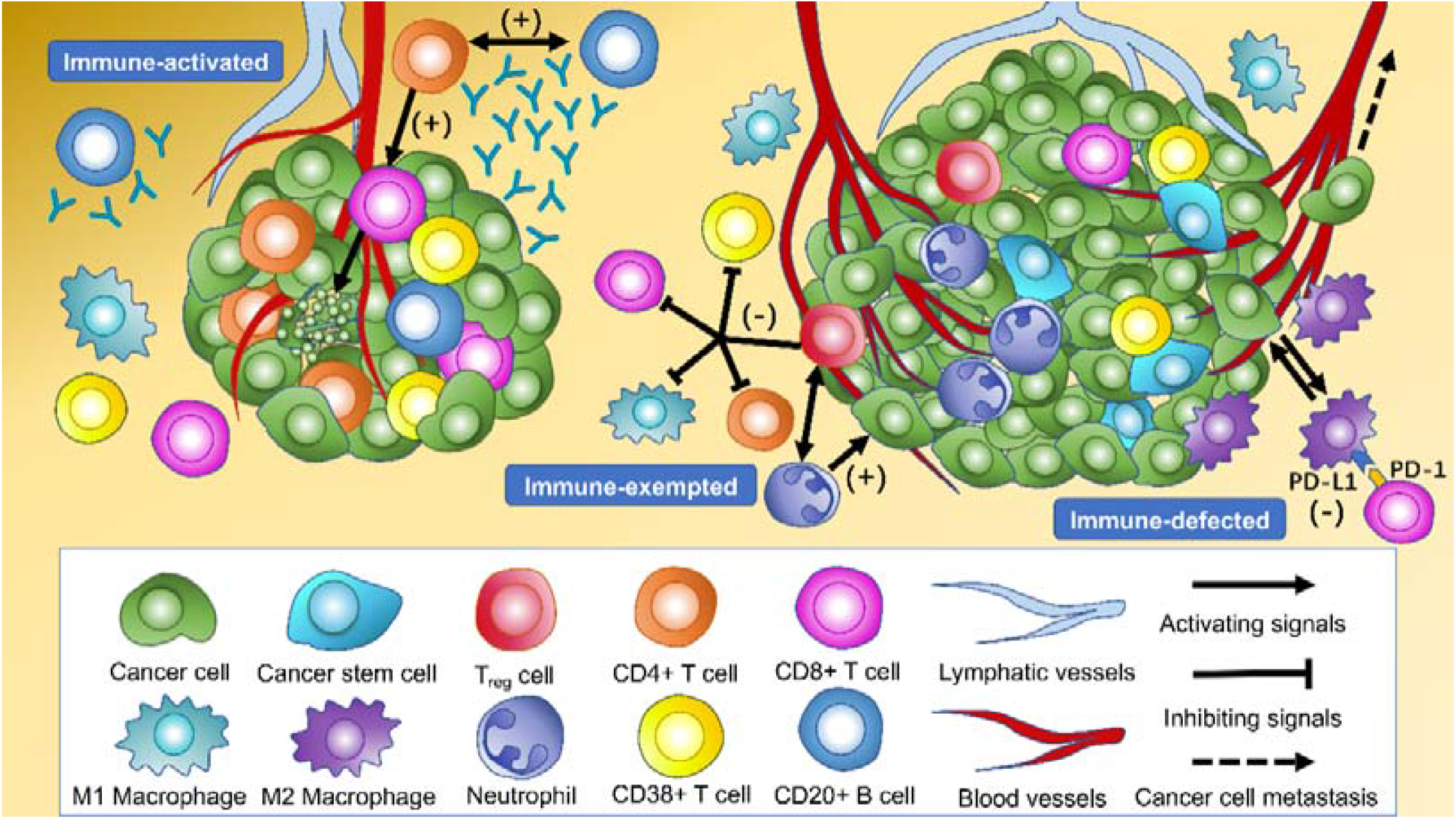
Cellular features of immune subtypes in tumor immune microenvironment (TIME) of non-small cell lung cancer (NSCLC). The immune-activated subtype is characterized by the highest levels of intratumoral and intrastromal CD4+ T cells, intrastromal CD20+ B cells, and intratumoral CD8+ T cells. CD20+ B cells present tumor antigens for activating CD4+ cells, and CD20+ B cells proliferate and differentiate into plasma cells for generating antibodies for antineoplastic effects with the help of cytokines secreted by CD4+ cells. CD8+ T cells activated by CD4+ cells tend to kill cancer cells in the tumor core directly. The highest levels of intratumoral and intrastromal regulatory T cells (Tregs) and neutrophils were observed in the immune-exempted subtype. Tregs produce immunosuppressive molecules which inhibit the activation and function of CD4+ T cells, CD8+ T cells, CD38+ T cells, and M1 macrophages to disrupt immune surveillance and promote tumor progression. Tregs may also recruit neutrophils through the chemokine ligand-chemokine receptor pathway. The immune-defected subtype has the highest levels of intratumoral and intrastromal cancer stem cells (CSC) and intratumoral macrophages. Macrophages in immune-defected subtype are educated by the CSC to obtain pro-tumorigenic functions like angiogenesis and induce the exhaustion of anti-tumor cells. The immune-exempted and immune-defected subtypes are associated with a more advanced-stage NSCLC than the immune-activated subtype.

### Prognoses of immune biomarkers

Twenty-eight out of 66 kinds of immune biomarkers were significantly associated with DFS. Intrastromal CD4-positive T cell was manifested as an independent protective biomarker in DFS. CD4++ T cells showed a stronger protective effect towards DFS than CD4+ T cells and CD4+++T cells. Moreover, intrastromal CD4+FOXP3+ Tregs were significantly associated with a longer DFS. CD8++T cells indicated the strongest protective effect towards DFS compared with CD8+T cells and CD8+++ T cells. Intratumoral CD8-positive T cells expressing CD133 exhibited strongest protective effect among all biomarkers (HR = 0.50, 95%CI 0.28-0.90). Immune cells expressing PD-L1 both in TN and TS were found to correlate with a longer DFS. Intrastromal neutrophils and CD20-positive B cells were related to the tendency of longer DFS.

Higher infiltration of macrophages was observed to be associated with worse outcome, and such effect was found to be more significant with the growth of fluorescence intensity. Further analysis revealed that both intrastromal and intratumoral CD68-positive macrophages expressing PD-L1 were associated with improved DFS, while CD68-positive macrophages without expressing PD-L1 were correlated with worse prognosis (Fig. 4). Moreover, a higher H-score (CD8) indicated a better DFS, whereas a higher H-score (CD68) demonstrated a worse DFS in TN.

**Figure 4.**
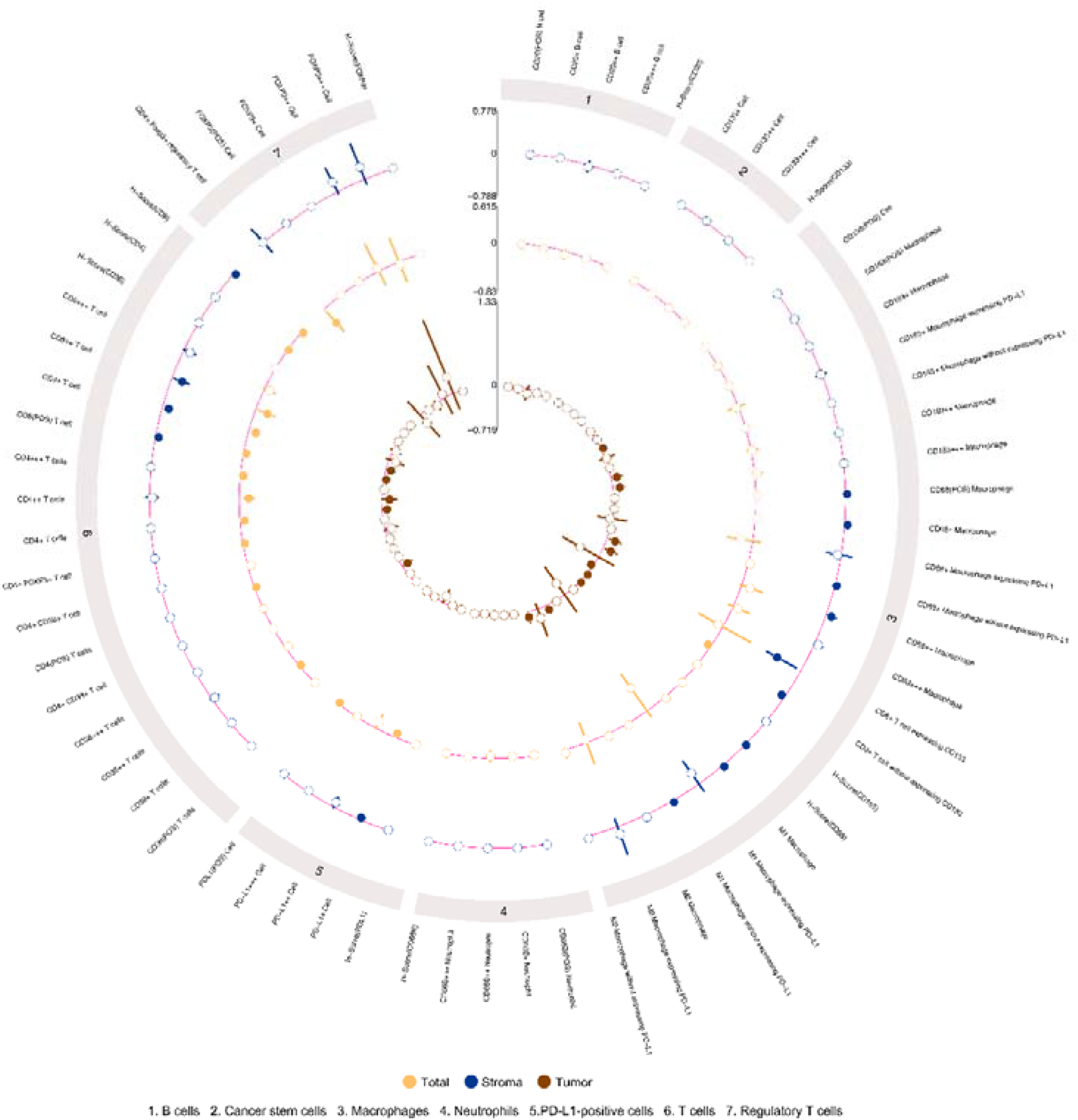
Circos plot demonstrates the prognostic significance of diverse immune biomarkers in tumor nest and tumor stroma as implement in multivariable Cox analysis with age, sex, T stage, N stage, vascular cancer embolus and number of lymph nodes resection as variates.

Higher H-score (CD4) and H-score (CD8) represented a better DFS in TS (Fig. 4, Supplementary Fig. S3). The prognostic effects of cell density and number of immune cells were available in Supplementary Table S6 and Supplementary Table S7. The distinction of infiltration levels of several immune biomarkers was statistically significant across T stage and N stage (Supplementary Fig. S4).

### Prognoses of immune-related risk score

Intratumoral CD68-positive macrophages, M1 macrophages and intrastromal CD4+ cells, CD4+FOXP3-cells, CD8+ cells, PD-L1+ cells were found to be the most robust prognostic biomarkers through LASSO (minimized lambda = 0.0281) and their regression coefficients derived for multivariable Cox proportional hazards regression analysis were 1.033, 1.035, 0.922, 0.968, 0.875 and 0.925, respectively (Fig. 5A, 5B). Hence, the following formula was utilized to calculate IRRS for each patient: IRRS = (intratumoral-%CD68-positive) * ln(1.033)+ (intratumoral-% M1 macrophages) * ln(1.035)+ (intrastromal-%CD4+) * ln(0.922) + (intrastromal-% CD4+FOXP3-cells) * ln(0.968) + (intrastromal-%CD8+) * ln(0.875) + (intrastromal-%CD8+) * ln(0.925). Multivariable Cox regression analysis showed that IRRS was significantly associated with DFS in the training cohort (p<0.001). We further stratified all patients into high-IRRS, medium-IRRS and low-IRRS subtypes using -0.01 and -0.86 as the optimal cutoff values. As illustrated in Table 1, the low-IRRS subtype had the most favorable DFS, whereas the high-IRRS subtype showed the worst (HR 2.63 95%CI 1.86-3.71, P<0.001), suggesting relatively great ability for risk stratification (Fig. 5C). Moreover, patients with median-IRRS also had longer DFS than patients with high-IRRS (HR 0.34, 95%CI 0.23-0.50, P <0.001). The prognostic performance of IRRS was assessed using time-dependent AUC curve, of which the training cohort indicated that the AUC fluctuated between 0.6-0.7, while AUC in the entire cohort gradually reach 0.9 after 2500 days, implying that the IRRS model had a higher predictive effect on the long-term risk in DFS (Fig. 5D, Supplementary Fig. S5). Moreover, with the increment of IRRS, the infiltrating levels of CD4-positive cells, CD8-positive cells, CD38-positive cells decreased gradually while the levels of macrophages increased gradually (Fig. 5E, 5F). Similar trends that patients in high-IRRS subgroup had significantly worse DFS than median-IRRS and low-IRRS subgroups were observed in the testing cohort (1031 days vs. 1710 days vs. 1792 days, P = 0.001) and the entire cohort (985 days vs. 1678 days vs. 1725 days, P < 0.001). The IRRS also showed potential ability for risk stratification (high vs. median vs. low) and prediction of five-year DFS rates (43.1% vs. 37.9% vs. 23.2%, P<0.001) in the entire cohort.

**Table 1.**
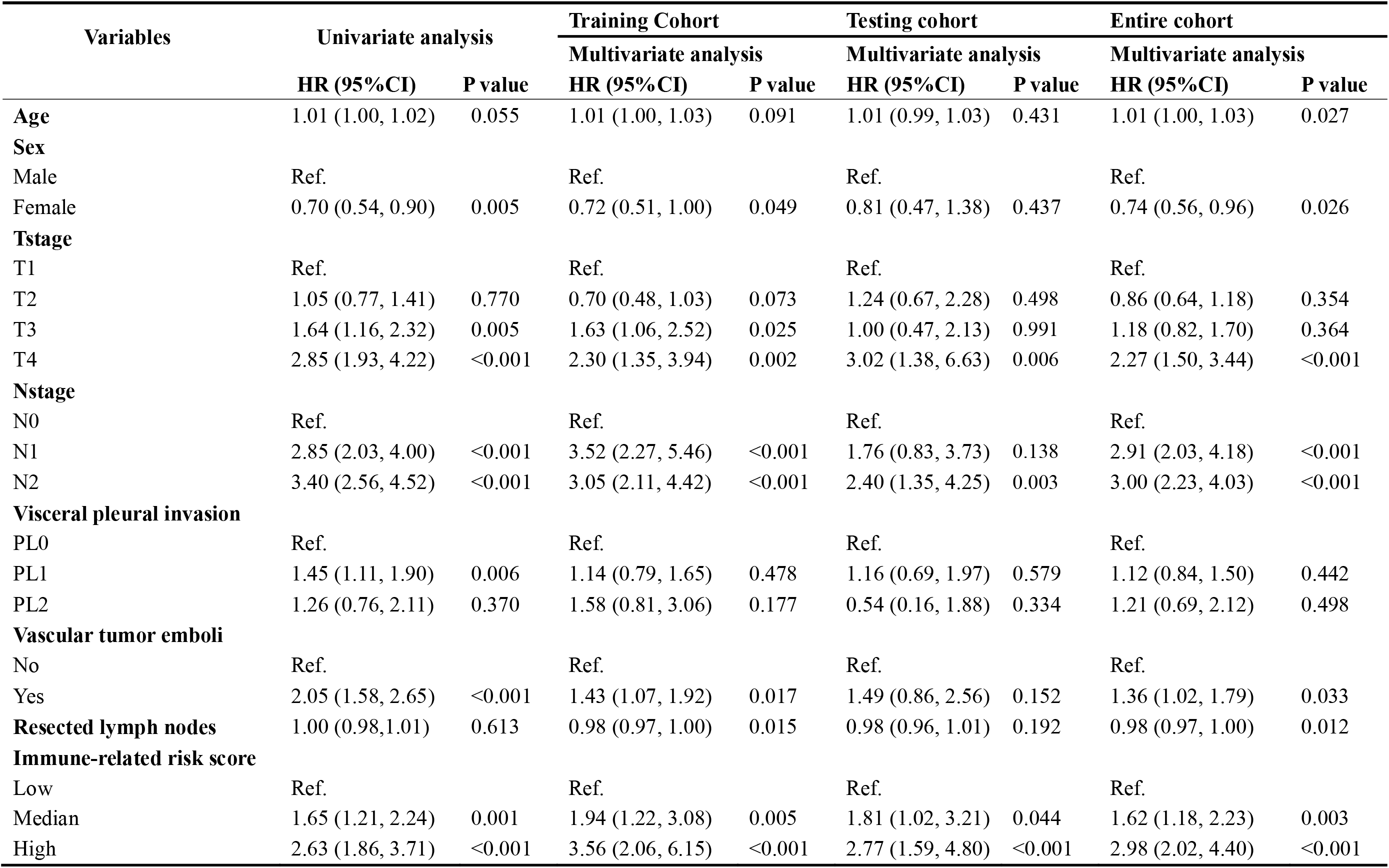
Construction and validation of immune-related risk score for predicting disease-free survival of non-small cell lung cancer.

| Variables | Univariate analysis |  | Training Cohort |  | Testing cohort |  | Entire cohort |  |
| --- | --- | --- | --- | --- | --- | --- | --- | --- |
|  | HR (95%CI) | P value | Multivariate analysis<br>HR (95%CI) | P value | Multivariate analysis<br>HR (95%CI) | P value | Multivariate analysis<br>HR (95%CI) | P value |
| **Age** | 1.01 (1.00, 1.02) | 0.055 | 1.01 (1.00, 1.03) | 0.091 | 1.01 (0.99, 1.03) | 0.431 | 1.01 (1.00, 1.03) | 0.027 |
| **Sex** |  |  |  |  |  |  |  |  |
| Male | Ref. |  | Ref. |  | Ref. |  | Ref. |  |
| Female | 0.70 (0.54, 0.90) | 0.005 | 0.72 (0.51, 1.00) | 0.049 | 0.81 (0.47, 1.38) | 0.437 | 0.74 (0.56, 0.96) | 0.026 |
| **Tstage** |  |  |  |  |  |  |  |  |
| T1 | Ref. |  | Ref. |  | Ref. |  | Ref. |  |
| T2 | 1.05 (0.77, 1.41) | 0.770 | 0.70 (0.48, 1.03) | 0.073 | 1.24 (0.67, 2.28) | 0.498 | 0.86 (0.64, 1.18) | 0.354 |
| T3 | 1.64 (1.16, 2.32) | 0.005 | 1.63 (1.06, 2.52) | 0.025 | 1.00 (0.47, 2.13) | 0.991 | 1.18 (0.82, 1.70) | 0.364 |
| T4 | 2.85 (1.93, 4.22) | <0.001 | 2.30 (1.35, 3.94) | 0.002 | 3.02 (1.38, 6.63) | 0.006 | 2.27 (1.50, 3.44) | <0.001 |
| **Nstage** |  |  |  |  |  |  |  |  |
| N0 | Ref. |  | Ref. |  | Ref. |  | Ref. |  |
| N1 | 2.85 (2.03, 4.00) | <0.001 | 3.52 (2.27, 5.46) | <0.001 | 1.76 (0.83, 3.73) | 0.138 | 2.91 (2.03, 4.18) | <0.001 |
| N2 | 3.40 (2.56, 4.52) | <0.001 | 3.05 (2.11, 4.42) | <0.001 | 2.40 (1.35, 4.25) | 0.003 | 3.00 (2.23, 4.03) | <0.001 |
| **Visceral pleural invasion** |  |  |  |  |  |  |  |  |
| PL0 | Ref. |  | Ref. |  | Ref. |  | Ref. |  |
| PL1 | 1.45 (1.11, 1.90) | 0.006 | 1.14 (0.79, 1.65) | 0.478 | 1.16 (0.69, 1.97) | 0.579 | 1.12 (0.84, 1.50) | 0.442 |
| PL2 | 1.26 (0.76, 2.11) | 0.370 | 1.58 (0.81, 3.06) | 0.177 | 0.54 (0.16, 1.88) | 0.334 | 1.21 (0.69, 2.12) | 0.498 |
| **Vascular tumor emboli** |  |  |  |  |  |  |  |  |
| No | Ref. |  | Ref. |  | Ref. |  | Ref. |  |
| Yes | 2.05 (1.58, 2.65) | <0.001 | 1.43 (1.07, 1.92) | 0.017 | 1.49 (0.86, 2.56) | 0.152 | 1.36 (1.02, 1.79) | 0.033 |
| **Resected lymph nodes** | 1.00 (0.98,1.01) | 0.613 | 0.98 (0.97, 1.00) | 0.015 | 0.98 (0.96, 1.01) | 0.192 | 0.98 (0.97, 1.00) | 0.012 |
| **Immune-related risk score** |  |  |  |  |  |  |  |  |
| Low | Ref. |  | Ref. |  | Ref. |  | Ref. |  |
| Median | 1.65 (1.21, 2.24) | 0.001 | 1.94 (1.22, 3.08) | 0.005 | 1.81 (1.02, 3.21) | 0.044 | 1.62 (1.18, 2.23) | 0.003 |
| High | 2.63 (1.86, 3.71) | <0.001 | 3.56 (2.06, 6.15) | <0.001 | 2.77 (1.59, 4.80) | <0.001 | 2.98 (2.02, 4.40) | <0.001 |

**Figure 5.**
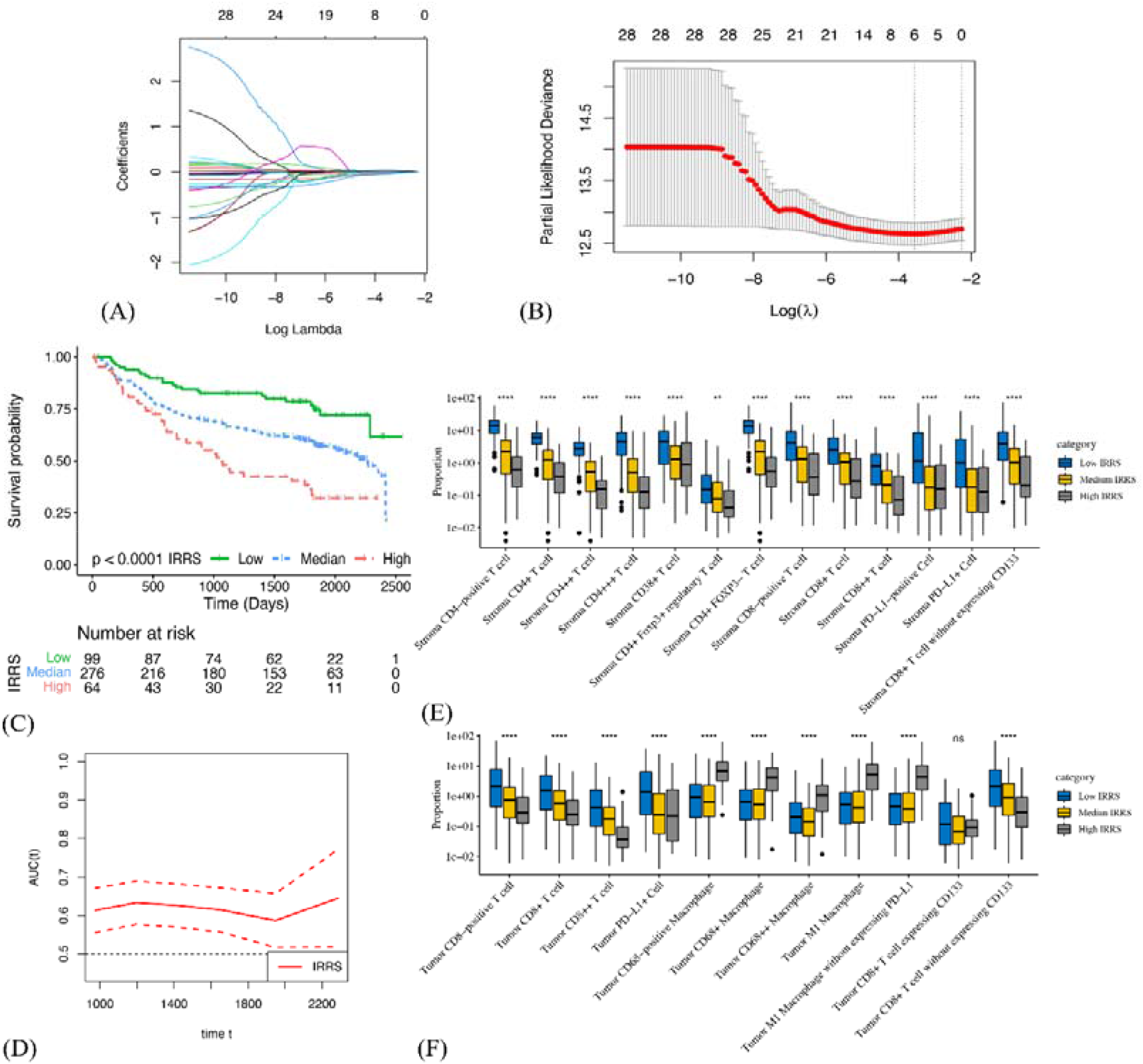
Construction and validation of immune-related risk score in the training cohort. (A) LASSO coefficient profiles of 28 selected immune cell biomarkers in the 10-fold cross validation. (B) Partial likelihood deviance revealed by the LASSO regression model in the 10-fold cross validation. (C) Kaplan–Meier estimate of the disease-free survival in the training cohort, divided by three IRRS subtypes. (D) Time-dependent AUC curve estimating the prognostic performance of IRRS in the training cohort. (E) and (F) Boxplots present the infiltration disparities of three IRRS subgroups in tumor nest and tumor stroma of the training cohort.

## Discussion

We presented a MIF method for simultaneous identification of colocalized biomarkers in immune cells phenotyping in TIME. This is the first study to highlight the comprehensive characteristics and clinical significance of in situ immune cells from resected NSCLC using a large cohort. Firstly, we identified three robust immune subtypes through unsupervised consensus clustering, including the immune-activated, immune-exempted, and immune-defected. Each of immune subtypes was correlated with distinct infiltrating immune cells levels, and accordingly indicating significantly different clinical outcomes. After that, we presented an IRRS utilizing multivariable Cox regression and LASSO-Cox regression analyses, clearly demonstrating the potential ability for risk stratification and prognosis prediction for DFS.

Our findings have several strengths and the following important aspects which differ from previous studies. This is the first study that investigated the density, proportion, number, and H-score for each immune cells in both tumor and paratumor stroma. We implemented the MIF test for 66 kinds of immune biomarkers from stage I to III in a large cohort of 681 patients, avoiding the shortcomings like homogeneous cohorts (a particular clinical stage), low statistical power, and wide variation., Moreover, except for using established biomarkers, we additionally tested other immune biomarkers like CD8+CD133+, CD4+CD38+ and further analyzed their functional orientation by the presence of PD-L1, reflecting multifarious immunological processes. In addition, traditional prognostic biomarkers were usually developed by an individual-based model which requires the information of clinical outcomes to be known in advance, namely “supervised”. On the contrary, we utilized unsupervised consensus cluster approach based on the levels of immune biomarkers profiles to reveal the intricate and intrinsic structure of TIME, maximizing the homogeneity of immune components within the same cluster and the heterogeneity between different clusters. Finally, we designed an IRRS based on quantitative evaluation of infiltrating immune cells specific to the constitution of TIME rather than non-specific gene signatures that were generally used in previous studies. The candidate factors were selected in a rigorous method based on multivariate Cox regression and LASSO-Cox regression analyses, enhancing the statistical power.

Study from Chen et al. which focused on head and neck cancer, presented three immune subtypes, namely non-immune, exhausted, and active, respectively (45). Similarly, the immune-activated class accounted for largest proportion of our patients. The immune-exempted subtype was consistent with Chen’s defined non-immune class which showed significantly lower infiltrating levels of lymphocytes and more advanced T stage. The immune-defected class, in which macrophage tended to exhibit pro-tumor activity under the impact of CSC, was, however, not been reported yet. Therefore, our findings recapitulated the immune classes and complemented previous studies. Noteworthily, intracluster heterogeneity was also observed in our analysis, suggesting novel methods for clustering should be developed in the future.

The impact of immune profiles in TIME on patients’ survival has been well described across cancer types. The immune-activated subtype in our study showed the highest infiltration of immune effectors like CD4+ T cells, CD20+ B cells, and CD8+ T cells without expressing CD133, and accordingly, patients in this class had the longest DFS. On contrary, immune-defected tumors had a mass of CSC and macrophages, indicating the worst outcome. Noteworthily, macrophages primarily originate from bone marrow and polarize by tumor-derived signals (46). Two major lineages, including M1 and M2 of polarization, have been well described. Generally, M1 macrophages mainly display antitumoral functions by secreting cytokines for T cell activation, while M2 macrophages are perceived as pro-tumor effectors through angiogenesis and chemotactic function of Tregs (47). Consistent with these findings, M2 macrophages without expressing PD-L1 were enriched in the immune-defected subtype.

However, we observed that higher infiltration of M1 macrophages was also associated with shorter DFS, and so the specific function of macrophages still needs to be evaluated synthetically. Immune-exempted tumors were dominated by neutrophils and Tregs, which were critical for creating an immunosuppressive microenvironment through TGF-β and IL-10 signaling, and accordingly, patients were in a more advanced clinical stage. Importantly, the role of neutrophils in LC was still divergent so far. Evgeniy B. and colleagues have previously reported neutrophils could stimulate T cell responses in early-stage LC by increasing IFN-γ production (42). In comparison, patients in immune-exempted were mainly enriched in advanced-stage, and so our findings may suggest that neutrophils tend to exhibit pro-tumor rather than anti-tumor effect in advanced LC.

Our findings may offer a reference for designing rational combination immunotherapy strategies. For instance, patients in immune-activated class may benefit from single-agent ICB, reinforcing the preexisting anti-tumor responses and further extending survival. As for immune-exempted and immune-defected subtypes, ICB alone may not be sufficient considering the presence of immunosuppressive mechanisms. In this regard, TGF-β inhibition (NCT02423343 and NCT04064190 are ongoing trials), radiotherapy, or chemotherapy plus ICB can be utilized to change non-inflamed malignancy into inflamed one and further stimulate the dampened anti-tumor immunity (48). Novel approaches for these two subtypes, like transferring of neoantigen-reactive T cells and NK cells which can enhance anti-tumor immunological effects, are under active investigation. For patients with intracluster heterogeneity, therapeutic selections should depend on the specific TIME, usage of targeting carcinoma-associated fibroblasts therapies (49), or anti-angiogenic therapies (50) plus ICB may work. In summary, further studies are in an urgent need to detect the exact molecular and cellular mechanisms responsible for immune inactivity for curating novel combination strategies.

Our study also provided evidence for complicated correlations among immune cells, like Tregs and neutrophils, CD20-positive B cells and CD4+CD38+ T cells, CSC and macrophages, implying the chemotactic function may contribute to the formation and evolution of TIME. Moreover, the prognostic significance of each immune biomarker, varying from low fluorescence intensity (+) to high fluorescence intensity (+++) were evaluated. Interestingly, we observed that the prognosis effects of median fluorescence intensity (++) were mostly more significant than low or high ones, deserving further investigation.

Several limitations existed in our study. Firstly, although we investigated the prognostic significance for as many kinds of immune biomarkers as possible, the biological mechanisms behind them were unclear, and further experimental studies are warranted. Secondly, our patient cohort did not include stage IV samples and the proportion of stage III samples were limited as well (0.5%). Therefore, further studies should pay more attention to covering the advanced-stage NSCLC. In addition, we could not make external validation of IRRS. Hence, the generalization of our findings needs to be confirmed by more studies. Finally, due to the lack of treatment information, we could not assess the value of IRRS in predicting treatment response.

In summary, we comprehensively demonstrated the immune landscape of NSCLC through MIF analysis and further identified three robust immune subtypes, which may help identify the ideal candidates and tailor rational immunotherapeutic strategies. We also revealed the prognostic significance of 66 kinds of immune biomarkers and subsequently constructed an IRRS for predicting patients’ DFS, attributing to the risk stratification and prognosis prediction for DFS. Future studies with a larger sample size and better design are warranted for our deeper understanding of TIME.

## Declaration

### Ethics approval and consent to participate

The authors are accountable for all aspects of the work in ensuring that questions related to the accuracy or integrity of any part of the work are appropriately investigated and resolved

## Consent for publication

All authors consent to publication

## Competing interests

The authors declare no potential conflicts of interest.

## Funding

This work was supported by China National Science Foundation (Grant number 81871893); Key Project of Guangzhou Scientific Research Project (Grant number 201804020030). Cultivation of Guangdong College Students’ Scientific and Technological Innovation (“Climbing Program” Special Funds) (Grant number pdjh2020a0480, pdjh2021a0407);

## Authors’ contributions

(I) Conception and design: H.X.P., X.R.W., and W.H.L.

(II) Administrative support: H.X.P., X.R.W., and R.Z.

(III) Provision of study materials or patients: T.Y., X.Y.C., J.L., Y.K.W., Y.Y.A., J.N.C., Y.T.L.

(IV) Collection and assembly of data: H.B.Z., Y.H.C.

(V) Data analysis and interpretation: H.X.P. and X.R.W.

(VI) Manuscript writing: All authors

(VII) Final approval of manuscript: All authors

## Acknowledgements

None.

## Abbreviations

AUC: Area Under Curve
CD: Cluster of differentiation
CDF: Cumulative distribution function
CI: Confidence interval
CTLA-4: Cytotoxic T-lymphocyte antigen 4
DCA: Decision curve analysis
DFS: Disease-free survival
FOXP3: Forkhead box P3
HR: Hazard ratio
H-score: Histochemical score
ICB: Immune checkpoint blockade
IFN-γ: Interferon-γ
IL: Interleukin
IRRS: Immune-related risk score
K-W: test Kruskal-Wallis test
LASSO: Least absolute shrinkage and selection operator
MMPs: Matrix metallopeptidases
NCCN: The National Comprehensive Cancer Network
NSCLC: Non-small cell lung cancer
OS: Overall survival
PCA: Principal component analysis
PD-L1: Programmed death-ligand 1
SCLC: Small cell lung cancer
TAMs: Tumor-associated macrophages
TGF-β: Transforming growth factor-β
TH: T-helper cell
TIL-Bs: Tumor-infiltrating B cells
TILs: Tumor-infiltrating lymphocytes
TIME: Tumor immune microenvironment
TLS: Tertiary lymphoid structure
Treg: Regulatory T cell
TSA: Tyramine signal amplification

## Supplementary materials

Supplementary Table S1. Immune biomarkers and corresponding immune cell types in multiplex immunofluorescence test.

Supplementary Table S2. Clinicopathologic characteristics of the included patients.

Supplementary Table S3. Correlation coefficients among various types of immune cells in our study.

Supplementary Table S4. P-values of correlation coefficients among various types of immune cells in our study.

Supplementary Table S5. Univariate and multivariate Cox regression analysis of disease-free survival, including immune subtype, age, sex, Tstage, Nstage, vascular tumor emboli, and number of resected lymph nodes of patients with non-small cell lung cancer.

Supplementary Table S6. The prognostic effects of number of immune cells evaluated by multivariable Cox regression analysis.

Supplementary Table S7. The prognostic effects of cell density of immune cells evaluated by multivariable Cox regression analysis.

Supplementary Fig. S1. Spearman’s rank correlation matrix (right half) and corresponding p-value (left half) among various intratumoral and intrastromal immune cell types. *, P<0.05, **, P<0.01, ***, P<0.001.

Supplementary Fig. S2. Identification of differences in spatial distribution of immune cells within tumor nest and tumor stroma using Kruskal-Wallis test. *, P<0.05, **, P<0.01, ***, P<0.001, “ns” means no significant difference.

Supplementary Fig. S3. Kaplan-Meier curves illustrate the association between disease-free survival and infiltrating proportion rate of immune biomarkers (high vs. low) within tumor nest and tumor stroma.

Supplementary Fig. S4. Several immune biomarkers significantly associated with disease-free survival (p<0.05) are selected and T-test is performed in identification of infiltration distinction across clinical stage, T stage and N stage in tumor nest (a) and tumor stroma (b).

Supplementary Fig. S5. Kaplan-Meier curves of three IRRS subgroups of the entire cohort (A). Time-dependent AUC curve estimating the prognostic performance of IRRS in the training cohort (B). Boxplots present the infiltration disparities of three IRRS subgroups in tumor nest (C) and tumor stroma (D) of the entire cohort.

Supplementary Fig. S6. Kaplan-Meier curves of three IRRS subgroups of the testing cohort (A). Time-dependent AUC curve estimating the prognostic performance of IRRS in the testing cohort (B). Boxplots present the infiltration disparities of three IRRS subgroups in tumor nest (C) and tumor stroma (D) of the testing cohort.

## Notes

**Conflict of interest statement** The authors declare no potential conflicts of interest.

### Competing Interest Statement

The authors have declared no competing interest.

